# Dynamics of Gut Microbiome, IgA Response and Plasma Metabolome in Development of Pediatric Celiac Disease

**DOI:** 10.1101/2020.02.29.971242

**Authors:** Khyati Girdhar, Qian Huang, Yusuf Dogus Dogru, Yi Yang, Vladimir Tolstikov, Martina Chrudinova, Amol Raisingani, Jonas F. Ludvigsson, Michael A. Kiebish, Noah W. Palm, Johnny Ludvigsson, Emrah Altindis

## Abstract

Celiac disease (CD) is an autoimmune disorder triggered by gluten consumption. To identify the role of gut microbes in CD onset, we performed a longitudinal study focusing on two important phases of gut microbiota development at ages 2.5 and 5 (n=16). We obtained samples from children who developed CD during or after the study (CD progressors) and age, sex, and HLA-matched healthy controls. CD progressors had a distinct gut microbiota composition and IgA-sequencing identified unique IgA targets in the gut. Three cytokines, one chemokine, and 19 plasma metabolites were significantly altered in CD progressors at age 5. Feeding C57BL/6J mice with taurodeoxycholic acid (TDCA), a 2-fold increased microbiota-derived metabolite in CD progressors, caused villous atrophy, increased intraepithelial lymphocytes (IELs), CD4+ T-cells, Natural Killer cells, and Qa-1 expression on T-cells while decreasing T-regulatory cells in IELs. Thus, TDCA drives inflammation in the small intestines that potentially contribute to the CD onset.

**Highlights:**

- CD progressors have a distinct gut microbiome composition compared to healthy controls in two important phases of gut microbiota development (age 2.5 and 5 years)
- CD progressors have more IgA-coated bacteria in their gut at age 5 compared to healthy controls. Further, IgA-sequencing identified unique bacterial targets in CD progressors.
- Three plasma proinflammatory cytokines and a chemokine were increased in CD progressors years before diagnosis, indicating an early inflammatory response.
- We identified 19 metabolites that are significantly altered in CD progress at age 5 and microbiota-derived TDCA increased two-fold.
- TDCA treatment in B6 mice increased CD4+ cells and NK cells while decreasing CD8+ T-regulatory (Treg) cells. It also increased Qa-1 expression on immune cells.

## Introduction

Celiac disease (CD) is a gluten-induced autoimmune disorder that is predicted to affect 1 in 100 individuals worldwide (Lebwohl et al., 2018). The adaptive autoimmune response in CD is characterized by gluten-specific CD4+ T cells, antibodies against gluten gliadin peptide, and the enzyme tissue transglutaminase (tTG) responsible for deamidating the gliadin peptide (Lebwohl et al., 2018; Lindfors et al., 2019). Almost all CD patients possess HLA-DQ2 or HLA-DQ8. However, although 20%-40% of the population in Europe and the USA carries these alleles, only 1% of individuals develop the disease (Wolters and Wijmenga, 2008). King et al. recently showed that the incidence of CD is increasing by 7.5% per year in the last decades (King et al., 2020). Furthermore, even among twins, the concordance of CD is not 100 % (Greco et al., 2002; Kuja-Halkola et al., 2016). These findings suggest that CD development is regulated by both genetic and environmental factors. Thus, understanding the environmental triggers of pediatric CD is important to develop alternative treatments to a gluten-free diet (GFD) and new tools to prevent CD.

Various environmental factors are implicated in the CD development (Lionetti and Catassi, 2015). Gut microbiome studies observed an altered microbial composition (Nistal et al., 2012) and fecal metabolite composition in both infant and adult CD patients (Cheng et al., 2013; Leonard et al., 2021; Olivares et al., 2018; Sellitto et al., 2012; Serena et al., 2017; Zafeiropoulou et al., 2020). However, previous studies have not identified any causal link between the gut microbiome and the disease onset or intestinal inflammation. In this study, we used fecal and plasma samples from a prospective, longitudinal cohort of 16 CD progressors and 16 healthy, age, HLA genotype, and breastfeeding duration-matched controls. To assess the gut microbiota and plasma metabolome alterations in pediatric CD onset, we used fecal samples obtained at ages 2.5 and 5 representing the two important stages of the gut microbiota development (Stewart et al., 2018). This analysis revealed that CD progressors have significant alterations in their gut microbiome composition, a unique IgA response against gut bacteria, and a distinct plasma metabolome and cytokine profile years before diagnosis. We showed that TDCA, a microbiota-derived metabolite that is two-fold increase in CD progressors, can exacerbate inflammation in the intestines in vivo. These findings open a new avenue for understanding the role of gut microbes and gut microbiota-related metabolites in CD onset.

## RESULTS

### Celiac Disease Progressors have a Distinct Gut Microbiota Composition

We first determined gut microbiome composition differences between CD progressors and healthy controls using 16S rRNA gene sequencing. The characteristics of the subjects are described in **Table S1**. In total, we identified 575 amplicon sequence variants (ASVs) (**Table S2**). Alpha and beta diversity were comparable between CD progressors and healthy subjects at age 2.5 and 5 (**Figure 1A**). Principal component analysis (PCA) showed a trend of separation of gut microbiome composition between CD progressors and healthy subjects at age 2.5 (**Figure 1B**). Relative abundance analyses of microbial taxa revealed that there were no significant differences among the phylum or genera levels (**Figure 1C and 1D, Table S3**). However, we identified significant differences at the ASV level in both ages. Specifically, 126 ASVs were different at age 2.5 and 109 ASVs were different at age 5, due to the small sample size (n=16), we used a very stringent analysis to report any significant finding (FDR <0.05, p <0.01; **Figure 1E**).

**Figure 1.**
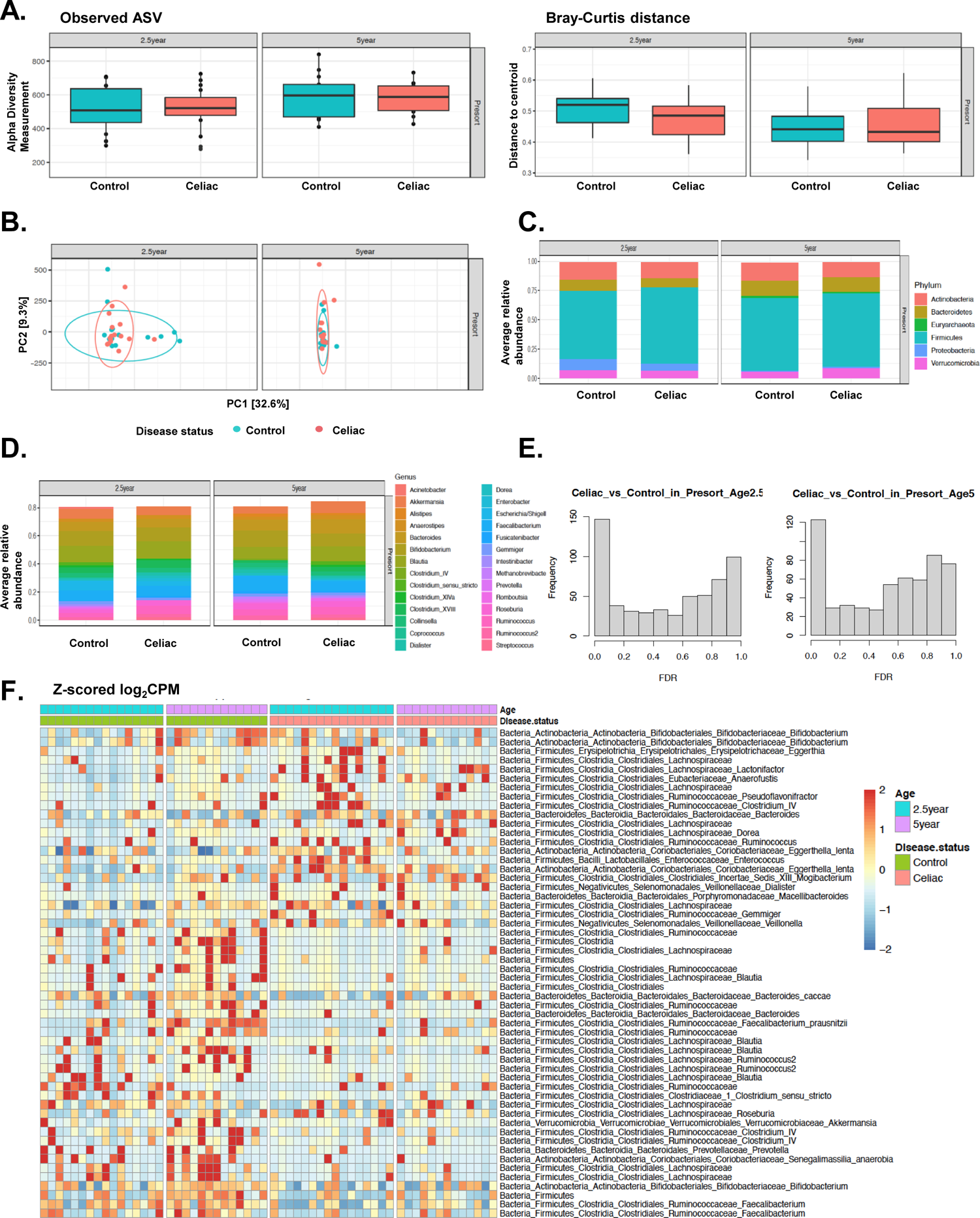
The Gut Microbiota of Children Developing CD and Healthy Controls were Different in the First 5 Years of Life. **A.** Box plots showing the comparison between CD progressors (n=13-16) and healthy controls (n=13-16): the alpha diversity measured by observed ASVs (left panel) and the beta diversity measured by Bray–Curtis dissimilarity (right panel). Statistical analysis was performed using ANOVA (alpha diversity) and PERMANOVA (Bray-Curtis distance). **B.** Principal component analysis (PCA) ordination of sample similarity/dissimilarity between CD progressors and healthy controls at age 2.5, and 5 years old. Each circle represents an individual sample, Control (green bar), Celiac (Red bar). **C-D.** Average relative abundance of bacterial phylum (upper panel) or genera (lower panel) of greater than 1% abundance (proportion) between the gut microbiota of CD progressors and healthy controls at age, 2.5, and 5 years old (taxa average relative abundance>1%). Statistical analysis was performed using two-tailed t-tests with Benjamini and Hochberg method to control False Discovery Rate (FDR). **E.** Empirical Bayes quasi-likelihood F-tests analysis for the comparisons of gut microbiota ASVs between CD progressors and healthy controls age 2.5, and 5 years old. Frequency: number of ASVs. **F.** Heat map showing the relative abundance of the top ASVs significantly different between CD progressors and healthy controls. Each column represents an individual participant and each row represents an ASV.

The most significantly enriched ASVs in CD progressors’ samples were *Dialister* (Fold Change (FC)=575), *Gemmiger* (FC=241), and *Clostridium_XVIII* (FC=108) at age 2.5. Likewise, the top species were *Dialister propionicifaciens* (FC=45.4)*, Phascolarctobacterium faecium* (FC=56.9), *and Porphyromonas asaccharolytica* (FC=19.6) at age 2.5 (**Figure S1A**). On the other hand, *Methanobrevibacter* (FC=817), *Clostridium sensu stricto* (FC=582), and in species-level *Bacteroides uniformis* (FC= 161), *Bacteroides stercoris* (FC=75.1), and *Parabacteroides goldsteinii* (FC=74.4) were highly enriched in healthy subjects (**Figure S1A**). Likewise, *P. faecium* (FC=419), *Dialister* (FC=1170), and *Ruminococcus* (FC=332) and *Dialister propionicifaciens* (FC=332), *Slackia piriformis* (FC=36.5) and *Shigella dysenteriae* (FC=14.2) were enriched in CD samples at age 5. Meanwhile, *Prevotella* (FC=-660), *Clostridium_IV* (FC=373) *Holdemanella* (FC=-387) and in species-level *Catenibacterium mitsuokai* (FC=237), *Bacteroides massiliensis* (FC=205), and *Bacteroides eggerthii* (FC=154) were enriched in healthy samples at age 5 (**Figure S1B**). The heat map shows a clear separation between CD progressors and healthy subjects at both ages (**Figure 1F**). These results indicate that the gut microbiota composition of CD progressors is significantly distinct in the first 5 years of life.

### CD progressors have More Bacteria Coated with IgA, Indicating an Inflammatory Gut Microbiota Composition

Immunoglobulin A (IgA) is the most abundant antibody isotype at mucosal surfaces and is a major mediator of intestinal immunity in humans (Bunker and Bendelac, 2018). IgA-sequencing (IgA-seq) combines bacterial cell sorting with high-throughput sequencing to identify distinct subsets of highly IgA coated (IgA+) and non-coated microbiota (IgA-) (Bunker et al., 2017; Palm et al., 2014; Planer et al., 2016; Wilmore et al., 2018). In this study, we used IgA-sequencing to determine the bacterial targets of the IgA in the gut microbiota of healthy children compared to CD progressors. The flow chart and the gating strategy are described in **Figures S2A and S2B**. PCA showed a separation between IgA+ and IgA-bacteria at all ages, both in control and CD samples (**Figure 2A**). The flow cytometry analysis revealed that there was a two-fold increase of the IgA+ bacteria in CD progressors (12.8%) compared to the controls (6.03%) at age 5 (p=0.026, **Figure 2B**). To determine whether this analysis is influenced by three children who developed CD before age 5, we removed these three samples and obtained a similar significant result (CD progressors: 12.8%, Control: 6.02%, p=0.027; **Figure S2C**). Overall, our data show that only a small fraction of the gut bacteria are coated by IgA in the first years of gut microbiota development in both healthy controls and CD progressors. However, CD progressors have more IgA+ coated bacteria at age 5 compared to healthy controls indicating an inflammatory gut environment for CD progressors years before the diagnosis.

**Figure 2.**
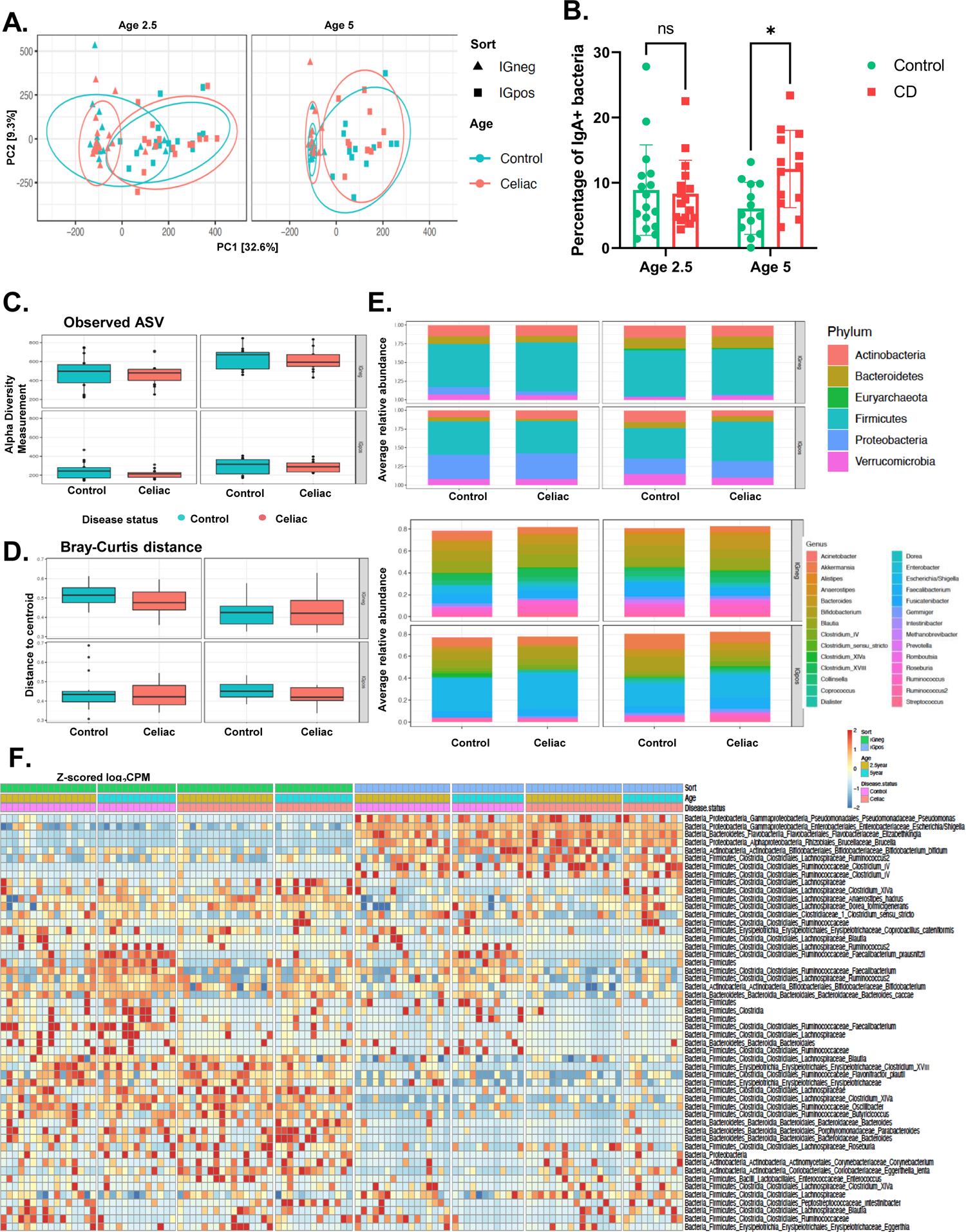
Gut microbiota of children developing celiac disease have more IgA+ bacteria than healthy controls at age 5. **A.** Principal component analysis (PCA) of sample similarity/dissimilarity between IgA+ and IgA-microbiota in healthy control (left) or CD progressors (right). **B.** Flow cytometry results for IgA + bacteria in fecal samples comparing CD progressors and healthy controls at age 2.5 and 5. Indicated are mean ± SEM. Data were expressed as means±SEM. *, P<0.05, **, P<0.01,***, P<0.001. Statistical analysis was performed by using two-way ANOVA. **C.** Box plots showing the comparison between CD progressors and healthy controls: the alpha diversity measured by observed for IgA+/IgA-microbiota at ages 2.5 (left panel), and 5 years old (right panel). Statistical analysis was performed using ANOVA. **D.** Box plots showing the comparison between CD progressors and healthy controls: the beta diversity measured by Bray–Curtis dissimilarity for IgA+/IgA-microbiota at ages 2.5 (left panel), and 5 years old (right panel). Statistical analysis was performed using PERMANOVA. **E.** Average relative abundance of IgA+/IgA-bacterial phylum (upper panel) or genera (lower panel) of greater than 1% abundance (proportion) between the gut microbiota of CD progressors and healthy controls at ages 2.5 (left panel), and 5 years old (right panel). Taxa average relative abundance>1%. Statistical analysis was performed using two-tailed t-tests with Benjamini and Hochberg method to control False Discovery Rate (FDR). **F.** Heat map showing the relative abundance of the top ASVs significantly different between IgA+ and IgA-samples of CD progressors and healthy controls (ASVs=51, selected based on p-value). Each column represents an individual participant and each row represents an ASV.

### A specific IgA response is already developed in both CD progressors and healthy children by age 2.5

There are very few reports on the IgA response in the early human gut microbiota development (Planer et al., 2016), thereby, we first focused on the results obtained from the healthy children. We identified 102 ASVs at age 2.5 and 41 ASVs at age 5 that were significantly different between IgA+ and IgA-samples in healthy controls (FDR<0.05, p<0.05; **Figure S2D**). The top targets of IgA in healthy subjects at age 2.5 were *Clostridium IV* (FC=3580) and *Bifidobacterium* (FC=190) and the top species were *Bacteroides clarus* (FC=140) and *C. mitsuokai* (FC=66.7). Likewise, *Clostridium IV* (FC=838), *Gemmiger* (FC=141), were the top enriched genera, and *B. clarus* (FC=64.9), and *Bifidobacterium bifidum* (FC=56.1) were the top enriched species at age 5 indicating an immunoregulatory role of these bacteria in healthy children.

On the other hand, we identified 112 ASVs at age 2.5 and 33 ASVs at age 5 that were significantly different between IgA+ and IgA-samples in CD progressors. These numbers are comparable to healthy subjects (FDR<0.05; **Figure S2D**). CD progressors shared similar top IgA targets with those in healthy subjects such as *Clostridium IV* (FC=1670), *Elizabethkingia* (FC=128) and *B. bifidum* (FC= 37.8) at ages 2.5 and *Clostridium IV* (FC=4020), *Gemmiger* (FC=346) and *B. clarus* (FC=97.8) at age 5. On the other hand, CD progressors’ targets of the IgA are *Lachnospiraceae* (FC=148), *Brucella* (FC=83.9), and at the species level, *Coprococcus comes* (FC=69), *Actinomyces turicensis* (FC=18.3) at age 2.5. *Enterobacter* (FC=514) *Clostridium XlVa* (FC=64.4), *B. clarus* (FC=97.8), and *Faecalibacterium prausnitzii* (FC=89.1) are the top targets at age 5 in CD progressors.

### IgA response targets are comprised of different bacteria in CD progressors

Consistent with the presorting data, both alpha and beta diversity were comparable in post-sorting samples (IgA -, IgA +) (**Figure 2C and 2D**). No difference was observed in the relative abundances of IgA+ or IgA-microbiotas at age 2.5 between CD progressors and healthy controls at the phylum level or genera level (**Figure 2E, Table S3**). On the other hand, we identified significant differences in the ASV level.

At age 2.5, IgA-Seq identified 131 different ASVs between control IgA-samples and CD IgA-samples. Likewise, we identified 155 different ASVs between control IgA+ samples and CD progressors’ IgA+ samples (FDR<0.05, **Figure S2E)**. At age 5, 69 ASVs were different between control IgA- and CD IgA-samples and 115 different ASVs were identified for CD IgA+ and control IgA+ samples (FDR<0.05, **Figure S2E**). The top differential targets of the IgA+ response enriched in CD progressors were *Clostridium IV* (FC=2480), *Gemmiger* (FC=137) and in species-level *Dialister propionicifaciens* (FC=238), *Bacteroides vulgatus* (FC=41.2) and *Clostridium sensu stricto* (FC=26.7) at age 5. Moreover, *Enterobacter* (FC= 458), *Clostridium XlVa* (FC= 305), *Roseburia* (FC=182), *Blautia* (FC=171), *Bacteroides* (FC=144), and in species-level *Dialister propionicifaciens* (FC=189), *Phascolarctobacterium faecium* (FC=307) and *Slackia piriformis* (FC=134) ASVs were highly coated with IgA in CD groups at age 5.

In addition to the differences caused by altered gut microbiota composition, we also identified 67 ASVs at age 2.5 and 46 ASVs at age 5 in which abundances were the same in the gut microbiota of CD and healthy samples (presorting) but differentially targeted by the immune system (**Table S4**). For example, *Bacteroides vulgatus (*FC=41.2), *Clostridium sensu stricto* (FC=26.7), *Clostridium XlVa* (FC=19.6) (**Figure S2F**), and *A. turicensis* (FC=14.4) ASVs at age 2. 5 and *Enterobacter* (FC=458), *Blautia* (FC=167), *Lactococcus* (FC=57.2,), and *Clostridium sensu stricto* (FC=24.8) ASV at age 5 were comparable in their abundance however they were highly coated with IgA in CD progressors but not in controls (**Figure 2F, Figure S2G**). Overall, these results indicate that gut microbiota composition and the IgA response to microbiota are distinct in CD progressors.

### CD progressors have increased inflammatory cytokines and chemokines years before the diagnosis

Our data showed that CD progressors have a distinct and stronger IgA response in the gut years before diagnosis. To determine if this is consistent with the systemic inflammation, we analyzed the cytokine profile at age 5 (**Figure 3A**). Assessing 48 cytokines (**Figure S3A)**, we identified three proinflammatory cytokines (IFNA2, IL-1a, IL-17E/(IL25)) and a chemokine (MIP-1b) that were significantly increased in CD progr essors years before diagnosis. In this analysis, we removed three children who were diagnosed with CD before age 5. IL-12p70, IL-27 also tended to increase (p= 0.051 and 0.058, respectively). Although our sample size was small, when we compared only CD patients to the healthy controls, we identified a significant increase in seven cytokines, three chemokines, and one growth factor that is potentially related to the disease (**Figure S3B**). This list includes all the significant alterations identified in CD progressors except IL-17E/(IL25). The IL-17E/(IL25) is specific to CD progressors and an important cytokine that was previously shown to play a key role in regulating autoimmune processes via T_H_17 cells (Borowczyk et al., 2021). Overall, the presence of these markers in CD progressors indicate that these children have increased inflammation years before diagnosis.

**Figure 3.**
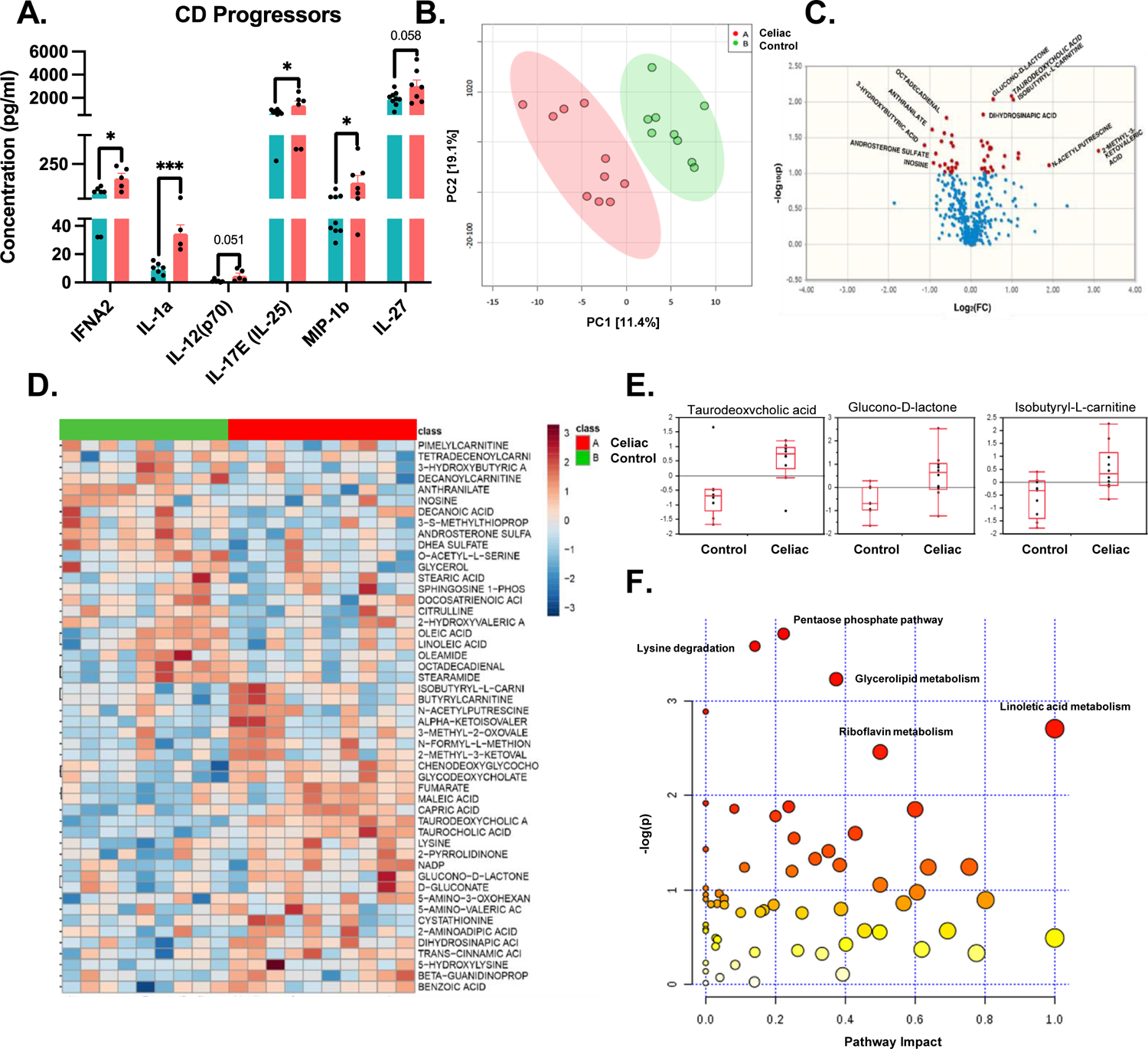
Plasma cytokines and metabolites levels alter in CD progressors. **A.** Cytokine profile of the CD progressors (n=7) in comparison to healthy controls (n=10) assessed by Luminex. Data were expressed as means ±SEM. *, P<0.05, **, P<0.01, ***, P<0.001. Statistical analysis was performed by a two-tailed, unpaired Student’s t-test. **B.** Partial Least Square-Discrimination Analysis (PLS-DA) of plasma metabolites for CD progressors (n=10) and controls (n=9). **C.** Volcano plot of plasma metabolites of CD progressors vs healthy controls with fold change threshold (|log_2_ (FC)|>1.2) and t-tests threshold (-log_10_(p)>0.1). The red dots represent metabolites above the threshold. Fold changes are log_2_ transformed and p values are log_10_ transformed. **D.** Heatmap showing 50 of the most altered metabolites between CD progressors and healthy controls. Each column represents an individual participant and each row represents a metabolite. **E.** Box plots showing three significantly altered metabolites in CD progressors. **F.** The Pathway Analysis (combined results from powerful pathway enrichment analysis with pathway topology analysis) identify the most altered metabolic pathways between CD progressors and healthy controls. Pathway impact value (x) is calculated from pathway topology analysis. p is the original p-value calculated from the enrichment analysis and depicted on a logarithmic scale.

### Plasma Metabolomics Analysis Reveals an Inflammatory Metabolic Profile for CD Progressors

In order to determine the early markers of CD progression in the plasma metabolome and its link to gut microbiota and inflammation, we applied a targeted plasma metabolomics analysis. We used plasma samples obtained at age 5 from CD progressors (including three-CD patients diagnosed before age 5) and healthy subjects. In total, we identified 387 metabolites, and partial least squares-discriminant analysis (PLS-DA) showed a clear separation of the plasma metabolites (**Figure 3B).** Volcano plots show the most significantly altered metabolites (**Figure 3C, Table S5**). Specifically, 19 out of 387 metabolites were significantly different (p < 0.05, **Table S5, Figure 3D**). The top three most altered metabolites (p < 0.01) were TDCA (FC=2), Glucono-D-lactone (FC=1.47), and Isobutyryl-L-carnitine (FC=2.058), and all three were increased in CD progressors (**Figure 3E**). In addition, the levels of two anti-inflammatory metabolites oleic acid (FC=0.69), and its derivative oleamide (FC=0.73), were significantly decreased in CD progressors (Oh et al., 2010). Interestingly, 2-Methyl-3-ketovaleric acid was increased more than eight folds in CD progressors (FC=8.6) (**Figure 3D**). Thus, we identified a strong plasma metabolome signature in CD progression (p<0.05).

One of the most altered metabolites, TDCA, is a conjugated bile acid that is mainly produced by gut microbes, particularly *Clostridium XIVa* and *Clostridium XI*, with 7-α-dehydroxylation of taurocholic acid and cholic acid (Ridlon et al., 2006). TDCA was previously shown to be a proinflammatory (Fujisaka et al., 2016). To determine whether this increase is influenced by the presence of three CD patients, we repeated our comparative analysis, comparing the metabolites of either CD patients or CD progressors with healthy controls, and observed a significant increase of TDCA in both groups independent of the diagnosis status (**Figure S3C and S3D**). Moreover, this result is consistent with our microbiota analysis since we identified several *Clostridium XIVa* ASVs that were significantly more abundant in CD samples, especially at age 5 (FC=39.5, FDR=0.01). Furthermore, *Clostridium XIVa* ASVs were highly targeted by IgA in CD progressors (FC=305, FDR=0.006) (**Figure S3E**). Using Pathway Analysis (**Figure 3F, Table S6**), we determined the functions related to these metabolites.

Consistent with the previous reports, we showed that the pentose phosphate pathway (PPP) (Raw P=0.0246) was significantly altered in the CD progressors (Leonard et al., 2021); in addition to lysine degradation (Raw P=0.028), and glycolipid metabolism (Raw P=0.0397) pathways.

### TDCA treatment resulted in villi atrophy and stimulated inflammation in C57BL6/J mice

To determine the effects of TDCA on the host, C57BL/6J male and female mice were fed either with standard chow or chow supplemented with TDCA (0.4% wt/wt) for ten weeks (**Figure 4A**). We did not observe any differences in the food intake caused by TDCA in the treatment group (**Figure 4B**). At the end of the treatment, we analyzed different cell subsets, including intraepithelial lymphocytes (IELs), lamina propria (LPs), mesenteric lymph nodes (MLNs), Peyer’s patches (PPs), and splenocytes. The MLNs of female mice fed with TDCA enlarged than those of the female control mice and tended to increase in male mice **(Figure 4C**). Further, both male and female mice fed with TDCA had a significantly higher number of cells in the MLNs indicating inflammation in the small intestines. TDCA treatment also decreased the number of intestinal epithelium cells (IECs) in both male and female mice (**Figure 4D).**

**Figure 4.**
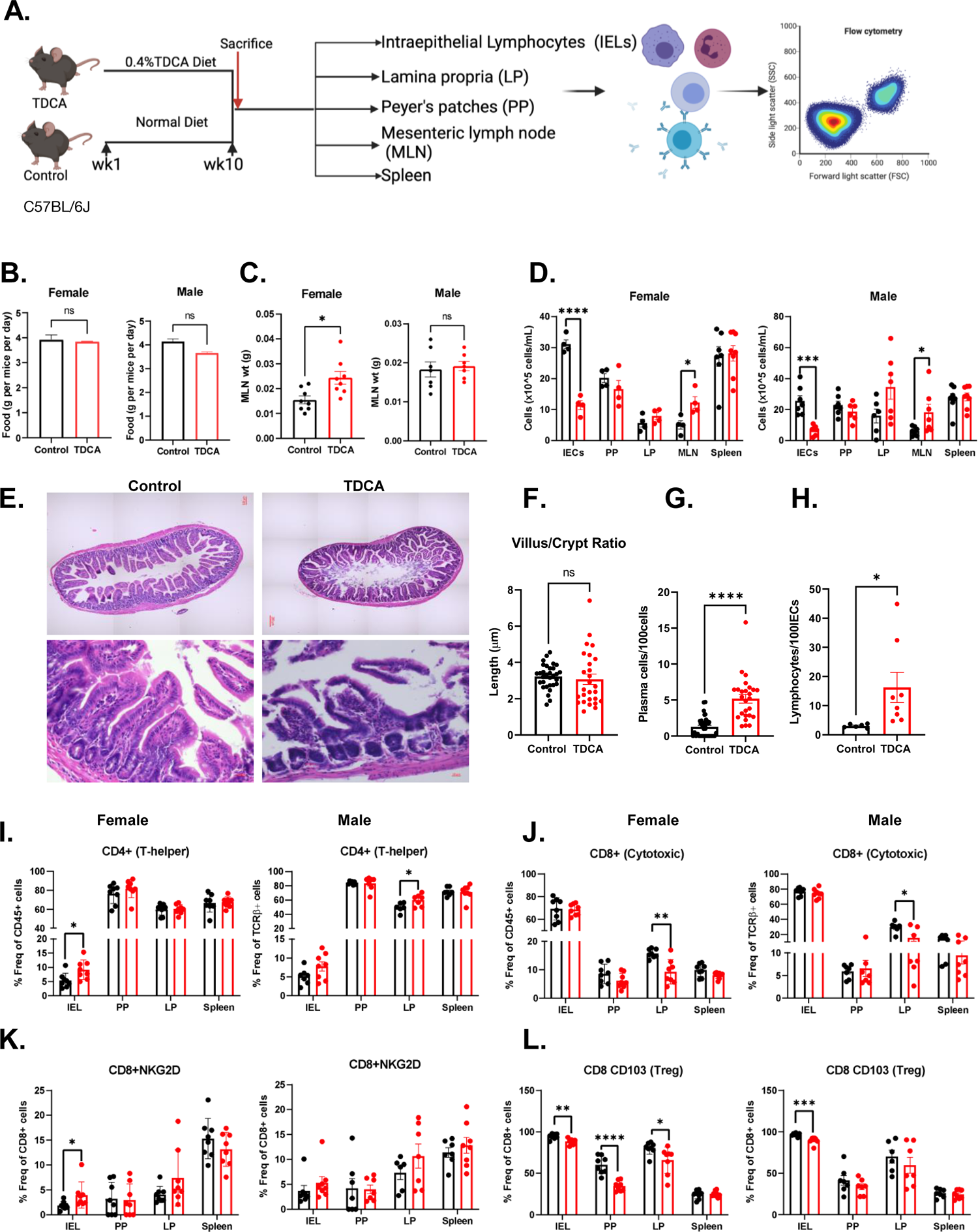
TDCA diet induces histological changes in the intestine and changes T-cell composition in different cell subsets. **A.** Schematic overview of the experiments (n=8/group/sex). Mice were fed either with a chow diet (control) or a chow diet containing 0.4% TDCA for 10 weeks. **B.** Food intake of mice represented as food intake per gram per mouse per cage. Female (left panel), male (Right panel). Control (black bar) and TDCA diet (Red bar) **C.** Weight of the mesenteric lymph nodes (MLN). Female (left panel) and male (right panel). **D.** Cell count was performed using a hemocytometer for each intestinal epithelium cell (IECs), Payer’s Patches (PP), Lamina Propria (LP), MLN, and spleen after isolation. Female (left panel) and male (right panel). **E.** H&E images of ileum tissue sections of control and TDCA treated female mice. Full image (upper panel) and image at high magnification (lower panel). Scale =20μm. **F.** Villi/ Crypt ratio in ileum tissue sections of control and the TDCA treated female mice. **G.** Number of plasma cells in the lamina propria of the ileum section of female mice. **H.** Number of IELs within 100 intestinal epithelium cells (IECs) in the villi of the female mice (ileum sections). **I.** CD4+ T-cells as % of total TCRβ+ cells in the IELs, PP, LP, and spleen of female (left panel) and male (right panel) mice. **J.** CD8+ T-cells as % of total TCRβ+ cells in the IELs, PP, LP, and spleen of female (left panel) and male (right panel) mice. **K.** CD8+ NKG2D+ T-cells as % of total CD8+ cells in the IELs, PP, LP, and spleen of female (left panel) and male (right panel) mice. **L.** CD8+ CD103+ T-cells as % of total CD8+ cells in the IELs, PP, LP, and spleen of female (left panel) and male (right panel) mice. Data were expressed as mean ± SEM. *P<0.05, **P<0.01,***P<0.01. Statistical analysis was performed by a two-tailed unpaired Student’s t-test.

The main characteristics of CD are an increase in intraepithelial lymphocyte (IELs), infiltration of plasma cells in the lamina propria, partial and total villus atrophy, and crypt hyperplasia (Dickson et al., 2006). To determine the histological changes in the intestines on treatment with TDCA, ileum sections of control and TDCA treated mice were stained with H&E. We observed distortion in crypt structure and partial and total villus atrophy caused by TDCA treatment in mice (**Figure 4E**). TDCA treatment didn’t cause any significant difference in the villi length, crypt depth, and villi/crypt ratio after 10wks of treatment (**Figure 4F**). However, we observed a five-fold increase in the plasma cells in the lamina propria (**Figure 4G**) and a 5-fold increase in the IELs in the intestinal epithelium cells (**Figure 4H**). These observations indicate that ten weeks of TDCA treatment was sufficient to cause villi atrophy and caused severe inflammation in the mice ileum.

### TDCA treatment increases CD4+ T-cells NKG2D receptor expression and decreases T-regulatory cells in-vivo

Next, we examined the different T-cell, natural killer (NK) cells population in IELs, PPs, LPs, and spleen after ten weeks of TDCA treatment. The gating strategy for the determination of all cell types has been described in **Figure S4**. TDCA-treatment induced a 2-fold increase in TCRβ+ cells in the PPs of both male and female mice (**Figure S5A**). Further, we identified a 2.1-fold increase in CD4+ T-cells in the IELs of TDCA-treated female mice and observed a similar trend for the male mice. There was also a 25% increase in CD4+ T-cells in LPs, specifically in male mice fed with TDCA (**Figure 4I**). TDCA treatment did not change the CD8+ T-cell population in IEL, PP, and spleen but decreased it in the LPs **(Figure 4J**).

NKG2D receptor is specifically expressed at the surface of all CD8+ αβT cells, γδT cells, and most NK cells (Wensveen et al., 2018) and plays a major role in the disruption of epithelium cells in CD pathogenesis interacting with the MHC class I chain-related proteins A (MICA) protein (Hue et al., 2004). MICA expression is increased under inflamed and stressed cells and serves as a ligand for NKG2D receptor for immune cell activation. TDCA treatment caused a 2-fold increase in the NKG2D receptor on T-cells in the IELs of the female mice, and the increasing trend was observed in male mice (**Figure S5B**). We then determined the NKG2D expression on CD8+ T-cells. TDCA treatment stimulated a 2-fold increase in the NKG2D expression in IEL of female mice and a trend of increase for the male mice (**Figure 4K**).

T-regulatory (Treg) cells play a key role in immune system homeostasis by immunosuppressing pathogenic T-cells (Gianfrani et al., 2006). Treg function is impaired in CD patients (Granzotto et al., 2009). CD8a+ TCRβ+ CD103+ cells, a novel subtype of CD8+ Treg cells, complement the function of CD4+ Foxp3+ Treg cells in suppression of immune response (Uss et al., 2006). We analyzed both CD4+CD103+ and CD8+CD103+ Tregs. TDCA treatment decreased CD4+CD103+ T-cells in IELs in both female (2.45-fold) and male mice (3-fold) (**Figure S5C**). TDCA treatment also decreased CD8+CD103+ cells by 15.7% in IELs, 42.5% in PP and 15.8% in LP as compared to female controls. We observed a decrease of 9.5% of CD8+CD103+ T-cells in IELs of male mice with TDCA treatment (**Figure 4L**).

TDCA treatment had a significant effect on the NK cell population in female mice and increased NK1.1+ cells in IEL (2.4-fold), PP (2.5-fold), LP (1.5-fold), and spleen (1.6-fold) (**Figure 5A**). Likewise, TDCA-treated male mice had an increase in the NK1.1+ cells in IEL (4-fold) and LP (2-fold) treatment (**Figure 5A**). It also increased NKp46+ cells in IEL (2.4-fold) and PP (3.3-fold) in female mice and IEL (2.4-fold) and LP (2-fold) in male mice (**Figure 5B**). Expression of two or more receptor activating receptors (such as NKG2D, NKp46, NKp44, DNAM1, NKp80, 2B4, and CD16) are required to trigger the activation of naïve NK cells. Thus, we analyzed both NKG2D+ NKp46+ markers on NK cells. Notably, the NKG2D+ NKp46+ population decreased by 1.3-fold in female splenocytes, by 2.7-fold in PP, and by 1.5-fold in splenocytes of male mice **(Figure 5C**). The decrease in these cells is relevant to CD onset because NKp44/NKp46 double-positive NK cells are significantly decreased in the duodenum of active CD patients (Marafini et al., 2016).

**Figure 5.**
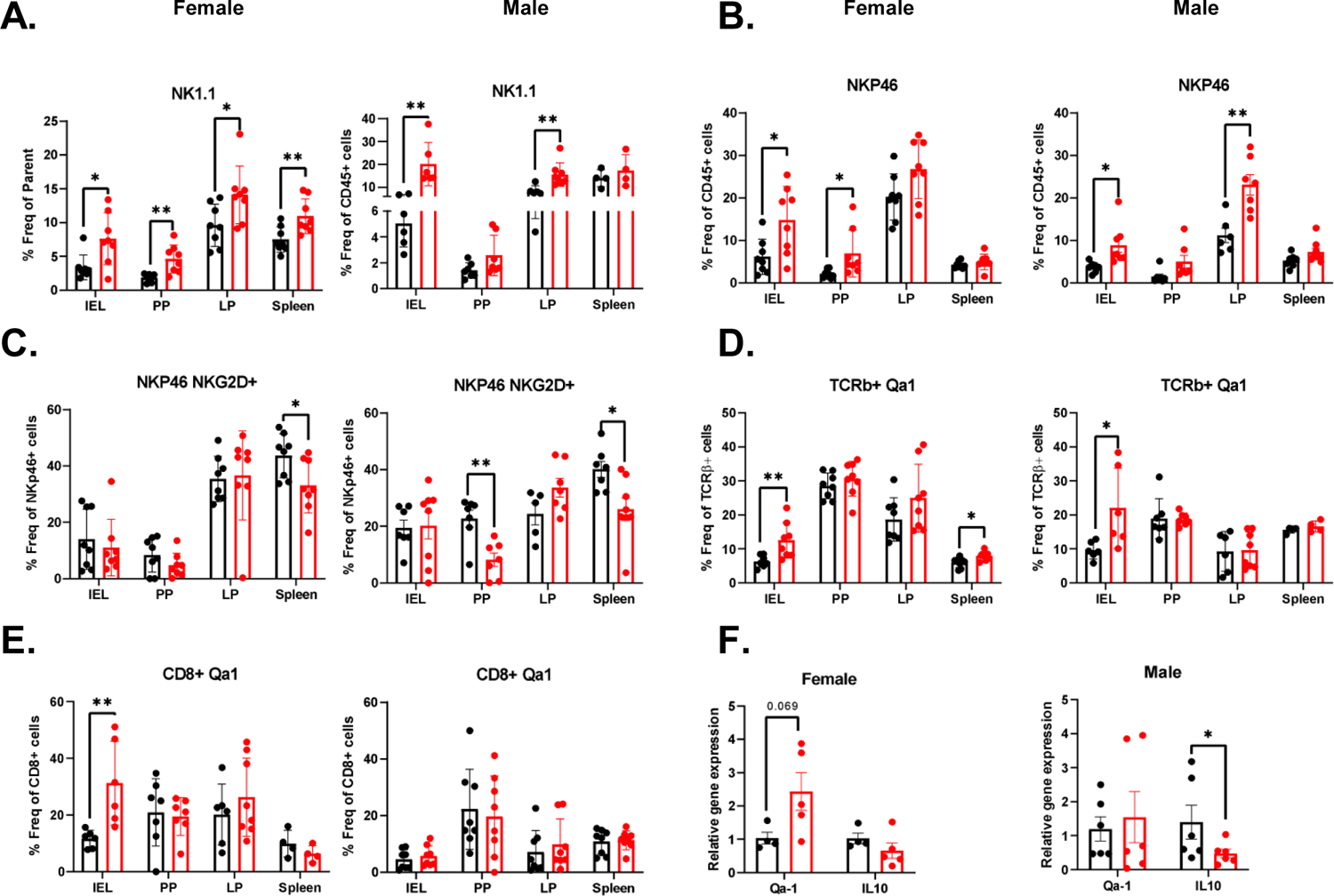
TDCA diet increases NK cells but decreases NK cells’ activity via increasing Qa-1 expression. **A.** NK1.1+ cells as % of total CD45+ cells in the IELs, PP, LP, and spleen of female (left panel) and male (right panel) mice. **B.** NKp46+ cells as % of total CD45+ cells in the IELs, PP, LP, and spleen of female (left panel) and male (right panel) mice. **C.** NKp46+ NKG2D+ cells as % of total NKP46+ cells in the IELs, PP, LP, and spleen of female (left panel) and male (right panel) mice. **D.** Qa-1+ cells as % of total TCRβ+ cells in the IELs, PP, LP, and spleen of female (left panel) and male (right panel) mice **E.** Qa-1+ cells as % of total CD8+ cells in the IELs, PP, LP, and spleen of female (left panel) and male (right panel) mice. **F.** Relative gene expression of Qa-1 and IL-10 in the ileum tissue analyzed using qPCR. Female (left panel), and male (right panel) mice after 10 weeks of TDCA treatment compared to controls. Data were expressed as mean ± SEM. *P<0.05, **P<0.01,***P<0.01. Statistical analysis was performed by a two-tailed unpaired Student’s t-test.

### TDCA treatment increases Qa-1 expression in cytotoxic T-cells, potentially protecting them from lysis

TDCA treatment stimulated a 2-fold increase in Qa-1, a murine homolog of HLA-E, expression in the TCRβ+ cells in IELs in both female and male mice, and a 1.3-fold increase in splenocytes of female mice (**Figure 5D**). It is also increased 3-fold on CD8+ T-cells of IELs in male mice **(Figure 5E**) and 1.2-fold on CD4+ T-cells of splenocytes in female mice **(Figure S5D)**. We also determined that the Qa-1 gene expression in the ileum tissue and Qa-1 tended to increase in the ileum of both female (p = 0.069) and male mice **(Figure 5F**). A previous study showed that the increased expression of Qa-1 on T-cells protects activated CD4+ T-cells from lysis by a subset of NK cells and disrupts the CD8+ Treg response (Lu et al., 2007). The decrease in Treg cell activity was observed in a reduction in IL-10 gene expression in the ileum tissue (**Figure 5F**. The reduction in the lysis of activated CD4+ T-cells results in more CD4+ T – cells, and defective Qa-1-restricted T-cell regulatory activity can result in autoimmunity (Kim and Cantor, 2011). Notably, in the Dd-V-IL-15-DQ8 mice model of CD, Qa-1 expression increases in the intestinal epithelium after gluten ingestion and is reduced when mice fed with gluten-free diet (Abadie et al., 2020), indicating an important link between Qa-1 and CD onset.

## DISCUSSION

Because most childhood CD cases will develop by age 5 years (Lionetti et al., 2014), we analyzed samples representing two critical gut microbiota developmental phases; a transitional phase and a stable phase (Stewart et al., 2018). Here, we report that CD progressors have a distinct microbiome, IgA response, and plasma metabolome and cytokine profiles compared to healthy children.

We identified significant differences in the ASV level, which is more informative about massive alterations in the phylum level. For example, the *Dialister* genus was one of the most abundant genera in CD progressors at both ages, and this is consistent with the previous reports (Zafeiropoulou et al., 2020). We observed a significant decrease in the abundance of some ASVs at age 2.5 that are the members of *Methanobrevibacter*, *Clostridium sensu stricto,* and *Bacteroides* (Sanchez et al., 2010) genera. These genera were previously reported to decrease in CD progressors and CD patients (Zafeiropoulou et al., 2020). Consistent with our data at age 2.5, it was previously shown that the abundance of several anti-inflammatory species, including *B. uniformis* (Leonard et al., 2021), *B. stercoris* (Cheng et al., 2013), and *Parabacteroides goldsteinii (Lai et al., 2022)* decreased in CD progressors as compared to healthy subjects.

Moreover, we identified ASVs from genera like *Prevotella*, that were decreased in CD progressors at age 5 and this genus was previously reported to decrease in CD patients (Di Biase et al., 2021). Likewise, we identified a decrease in ASVs identified in *Holdemanella* genus and it was previously reported that *Holdemanella* genus decreases in untreated pediatric CD patients, and treatment with GFD increases its abundance (Zafeiropoulou et al., 2020).

Planer et al. described mucosal IgA response progression during two postnatal years in healthy US twins (Planer et al., 2016). We used a similar approach to investigate the gut IgA immune development towards healthy and CD states; we initially focused on the development of the IgA response during the gut microbiota maturation. Flow cytometry analyses showed that the IgA response is highly selective, and only a small fraction of the gut microbiota is highly coated with IgA in the first five years of life. More importantly, the percentage of IgA+ bacteria were higher in CD progressors compared with healthy controls at age 5. While a reduction of secretory IgA (sIgA) in infant (4-6 months) fecal samples in CD progressors (Olivares et al., 2018) was reported previously, we did not observe such a defect in our cohort; in contrast, we identified a two-fold increase in the number of IgA coated bacteria in CD progressors, especially at age 5, however it is important to note that the sampling ages are very different in these two cohorts.

Although the abundance *Bacteroides vulgatus*, *Clostridium sensu stricto, Clostridium XlVa*, and *A. turicensis* ASVs were comparable between groups, they were explicitly targeted in the CD progressors at age 2. 5 and *Enterobacter*, *Blautia, Lactococcus,* and *Clostridium sensu stricto* were targeted at age 5. The abundance analysis shows there were no significant differences at genus levels between CD progressors and healthy subjects.

*Bifidobacterium* species can stimulate the production of IgA in the intestines (Fukushima et al., 1998; Park et al., 2002) and are known to regulate immune response by inducing dendritic cells and Treg cells with regulatory activities (Konieczna et al., 2012). Treg cells play an essential role in maintaining tolerance to food antigens and microbiota to suppressing autoimmunity and inflammation in the gut (O’Mahony et al., 2008; Sun et al., 2007). In our study, we showed that *Bifidobacterium bifidum* is targeted by the IgA specifically in the healthy children at age 5. Likewise we identified an ASV, *Bifidobacterium* that is targeted in healthy controls at age 2.5. These results suggest *Bifidobacterium* species have an immunoregulatory function in healthy individuals.

Our analysis revealed 155 ASVs at age 2.5 and 115 ASVs at age 5 that were highly IgA coated in the CD progressors. Among them, *Coprococcus comes*, *Lachnospiraceae* at age 2.5, and *F. prausnitzii* and *Clostridium_XlVa* at age 5 were the top targets of the mucosal immune response in CD progressors. Among these bacteria, *C. comes* been recently identified as the main IgA target in the human colon (Sterlin et al., 2019). Notably, we identified 67 ASVs at age 2.5 and 46 ASVs at age 5 that are equally abundant in CD progressors and healthy controls but selectively targeted by IgA in CD progressors. For example, *A. turicensis*, Clostridium XlVa, and *Bacteroides vulgatus* at age 2.5 and *Lactococcus* and *Blautia* at age 5 were selectively targeted by IgA in CD progressors but not in controls. *B. vulgatus* and *A. turicensis* were implicated in the development of gut inflammation, and a previous report identified this bacterium as enriched in CD patients (Li et al., 2018). It was previously shown in a mice model that IgA+ microbiota could induce more severe colitis than IgA-microbiota (Palm et al., 2014). Similarly, IgA-seq identified *Escherichia coli* as an inflammatory bacterium enriched in Crohn’s disease-patients with spondyloarthritis (Viladomiu et al., 2017). Further studies are needed to determine the role of IgA+ bacteria in CD pathogenesis.

Although the hallmark of CD is an intestinal inflammation, the disease affects different tissues. To determine the systemic effects, we measured the levels of 48 different cytokines, chemokines, and growth factors. Five proinflammatory cytokines (IFNA2, IL-1a, IL-12p70, IL-27, IL-17E/(IL25)) and one chemokine (MIP-1b) were increased in the CD progressors years before the diagnosis. IL-12, IL1a, and IFNA2 cytokines were previously shown to increase in CD patients and a decrease with the GFD (Manavalan et al., 2010; Monteleone et al., 2001). IL-12 (p70) has also been shown to increase in children with CD in a former study (Bjorck et al., 2015). Likewise, consistent with our results, MIP-1b/CCl4 chemokine increased in non-treated celiac disease patients (Bjorck et al., 2015). Among these, IL-17E (IL-25) was the only cytokine identified in CD progressors but not in the CD patients. This “barrier cytokine,” is interesting because it was shown to play a key role in regulating autoimmune processes via T_H_17 cells and maintain homeostasis by alarming the immune cells about the tissue injury (Borowczyk et al., 2021). Overall, our findings indicate the presence of early inflammation years before the diagnosis of CD in these children.

Our analysis revealed that plasma metabolites were significantly altered prior to diagnosis in CD progressors. The most significantly altered plasma metabolites were TDCA and isobutyryl-L-carnitine (**Figure 4C**), in which both were increased two-fold in CD progressors. Pathway analysis for plasma metabolites identified several pathways, including the pentose phosphate pathway (PPP), lysine degradation, and glycerolipid metabolism. A recent study on fecal metabolome in CD progressors showed that PPP is one of the most significantly increased pathways (Leonard et al., 2021). PPP is the most significantly altered pathway in our analysis.

This pathway stimulates the formation of NADPH as an antioxidant by regenerating reduced glutathione and neutralizing the reactive oxygen intermediates (ROI), thereby controlling cell inflammation (Perl et al., 2011). Thus, our results show that the plasma metabolites of CD progressors are distinct and potentially a component of the inflammatory response.

TDCA is a conjugated bile acid shown to be a proinflammatory (Fujisaka et al., 2016) and is mainly produced by gut bacteria, particularly by *Clostridium XIVa* and *Clostridium XI* (Ridlon et al., 2006). This observation suggests that the plasma TDCA detected in our study is secondary to some *Clostridium XIVa* species’ increased abundance in CD progressors at age 5. In this study, we showed that chronic TDCA exposure causes villous atrophy in C57B6/J mice after ten weeks of treatment while increasing IELs in the intestinal epithelium, plasma cells in the lamina propria, and CD4+ T-cells in IEL of the female and LP of the male mice. These are the main phenotypic changes reported during the CD progression (Lauwers et al., 2015). Thus our findings link a microbiota-derived metabolite, TDCA, to the CD progression related alterations in the small intestines.

In CD, CD4+ T-cells secrete several cytokines, increase T-cell expansion, and participate in the killing of intestinal epithelial cells via IELs (Jabri and Sollid, 2017). Previous studies have shown that T-cell mediated killing of intestinal epithelium cells in CD are regulated by NKG2D receptor expression on a set of immune cells and NKG2D receptor’s interaction with MICA protein (Hue et al., 2004). NKG2D is mainly expressed on CD8+ αβT-cells, γδ-cells, and NK cells where MICA expression increases in intestinal epithelium cells in the presence of cellular distress caused by gliadin in the CD (Martin-Pagola et al., 2004). Here, we show that TDCA treatment induces the expression of NKG2D on T-cells and on CD8+ αβT-cells in IELs of female mice. In addition to this increase in NKG2D, we also identified a decrease in CD8+ Treg cells. CD8+ Treg cells lyse effector T-cells via perforin and other cytokines (Wang and Alexander, 2009). Thus, TDCA stimulated a decrease in CD8+ Treg cells, potentially contributing to increased inflammation via increased effector T-cells in the small intestines of CD progressors.

NK cells express several activating and inhibitory receptors to regulate their cytotoxic activity. NKG2A serves as the inhibitory receptor while NKG2D, NKp44, NKp46, NKG2C, and NKp30 are the activating receptors. In our study, we show that TDCA treatment reduces NK cell activation by downregulating the surface expression of NKG2D/NKp46 receptors in splenocytes and both male and female mice and PP of the male mice. This is also relevant to the CD onset because NKp44/NKp46 double-positive cells, which are responsible for the enhanced production of granzyme B, are reduced in active CD patients (Marafini et al., 2016).

A subset of CD8+ Treg cells recognizes Qa-1, which is essential for the maintenance of self-tolerance (Choi et al., 2020). Qa-1 inhibits NK cells-mediated T-cell killing via suppressing CD8+CD103+ Treg-cells. Engagement of Qa-1 with the NKG2A receptor induces inhibitory signals for CD8+ Treg cells as well as NK cells and this can increase autoimmune disease risk as shown for experimental autoimmune encephalomyelitis (Hu et al., 2004). Here, we observed an increase in Qa-1 expression, a decrease in CD8+ Treg cells, and suppression of NK cell activation altogether resulting in accelerated inflammation in the small intestines of the TDCA-treated mice.

Currently, the only way to treat CD is strict adherence to a GFD, but 20% of patients do not respond to GFD and continue to have persistent or recurrent symptoms (Stasi et al., 2016). CD permanently reshapes intestinal immunity, and alterations of TCRγδ+ intraepithelial lymphocytes, in particular, may underlie non-responsiveness to the GFD (Mayassi et al., 2019). Our findings suggest that the inflammatory nature of the CD progressors’ gut microbiota, especially the increased TDCA, is potentially one of the early key components of intestinal inflammation in CD. The pro-inflammatory factors identified in this study have the potential to trigger local and systemic inflammation independent of the diet and may explain a failure to respond to GFD in some patients.

The main strength of this study lies in the longitudinal sampling that represents two important phases of gut microbiota development in children. Further, applying IgA-seq analysis provides an important dimension to evaluate the gut microbiome and the mucosal immune response in CD pathogenesis. The plasma metabolome analysis and identification of an increase in the inflammatory microbial metabolite, TDCA, is another important finding. Further, testing the effects of TDCA exposure in vivo identified a new link between intestinal inflammation and gut microbiota in CD. We used 16S sequencing as other groups did in the CD context (Bibbo et al., 2020; Leonard et al., 2021; Olivares et al., 2018) although this is a limitation compared to a shotgun sequencing approach, we were able to identify several ASVs in the species level.

Taken together, our findings identified a distinct gut microbiota, IgA response, plasma metabolome, and cytokine profile for the CD progressors years before diagnosis. Further, we establish a potential link between TDCA, a metabolite specifically produced by the gut microbes, and intestinal inflammation. TDCA has potential to serve as an early diagnostic marker of the disease; furthermore, targeting TDCA-producing bacteria early in life could be a preventative tool. These microbes/compounds may also complement a gluten-free diet in patients that continue to experience persistent CD symptoms. Understanding the role of the gut microbiota in CD onset will open novel avenues to understand disease pathogenesis and reveal new preventive and treatment models.

## Abbreviations

ABIS: All Babies in Southeast Sweden

APCs: Antigen-presenting cells

CD: Celiac disease

CDP: Celiac disease progressors

FC: Fold change

FDR: False discovery rate

GF: Gluten free

HLA: Human leukocyte antigen

ICD: International classification of disease

IgA: Immunoglobulin A

IgA-seq: Immunoglobulin A sequencing

KEGG: Kyoto encyclopedia of genes and genomes

MAA: Mean average abundant

MACS: Magnetic activated cell separation

LS: Least squares

NMDS: Non-metric multidimensional scaling

ASV: Amplicon Sequence Variant

PCA: Principle component analysis

PICRUSt: Phylogenetic investigation of communities by reconstruction of unobserved states

PLS-DA: Partial Least Square-Discrimination Analysis

PPP: Pentose phosphate pathway

sIgA: Secretory IgA

SNDR: Swedish National Diagnosis Register

TDCA: Taurodeoxycholic acid

tTG: Tissue transglutaminase

PP: Peyer’s patches

MLN: Mesenteric Lymph node

LP: Lamina propria

IEC: Intestinal epithelium cells

IEL: Intraepithelial lymphocyte

MICA: MHC class I polypeptide-related sequence A

NK: Natural Killer

CD: Cluster of Differentiation

NKp46: Natural killer cell p46-related protein

NKG2: Natural killer group 2

MCP-3: Monocyte-chemotactic protein 3

MIP-1b: Macrophage Inflammatory Protein-1 beta

Th1: T-helper type 1

IFNA2: Interferon Alpha 2

GROa: Growth-regulated alpha protein

FGF2: Fibroblast Growth Factor 2

IL: Interleukin

## Declarations

### Ethics approval and consent to participate

The study was approved by the Research Ethical Committees of the Faculty of Health Science at Linköping University, Sweden, (Dnr Linköping 287-96, Linköping 03-092, and Linköping 2018/380-32) and the Medical Faculty of Lund University, Sweden (Dnr Lund 83-97). In addition to video film presentation, oral and written informed consents were obtained from the parents of the children included in the study.

### Consent for publication

Not applicable

### Availability of data and materials

All ASV-related data analyzed in this study are included in this published article (Additional file 2: Tables S2). The 16S rRNA gene sequencing raw data generated in this study is available through the NCBI Sequence Read Archive Bioproject PRJNA631001. The plasma metabolomics data are included in this published article (See Additional file 14: **Table S8**). The gut microbiome analysis codes generated in this study are available at this link: https://github.com/altindislab/celiac-gut-microbiome.

## Author contributions

K.G and E.A designed research and wrote the paper. Y.D.D assisted with the bioinformatics analysis. K.G, A.R. assisted with all animal experiments, FACS staining and analysis while M.C and E.A assisted with some of the animal experiments. J.L is the Head of the ABIS study and assisted with human fecal sample collection, classification, and interpreting the data. Q.H, Y.Y, and N.W.P, assisted with IgA-seq experiments and analysis. V.T and M.A.K assisted with plasma metabolomics analysis. J.F.L assisted with research design and writing. All authors helped the analysis of the data that they contributed to producing and approved the final version of the manuscript.

## Acknowledgments

We are grateful to all children participating in the ABIS study, and their parents. Thanks also to Ingela Johansson, KEF, Linköping, for her skillful work with the samples, and Åshild Faresjö for registering data. The authors also want to thank Hui Pan, Jonathan Dreyfuss (Joslin Diabetes Center Bioinformatics Core) for their help with bioinformatic analysis and statistics. The authors would also like to acknowledge Patrick Autissier for the cytometry service (Flow Cytometry Core of Boston College), Bret Judson (Microscopy core of Boston Collge). Thanks to Danielle Stephens and Alpdogan Kantarci (Multiplex Core, Forsyth Institute) for their support with the multiplex cytokine analysis. We also thank BC undergraduate students Typhania Zanou, Yena Sung, Kaan Sevgi, Maximilian Figura, and Jaewon Oh and graduate rotation students, Minqi Shen, Jelena Momirov and Kristina Kelley for their technical help.

## Additional Files

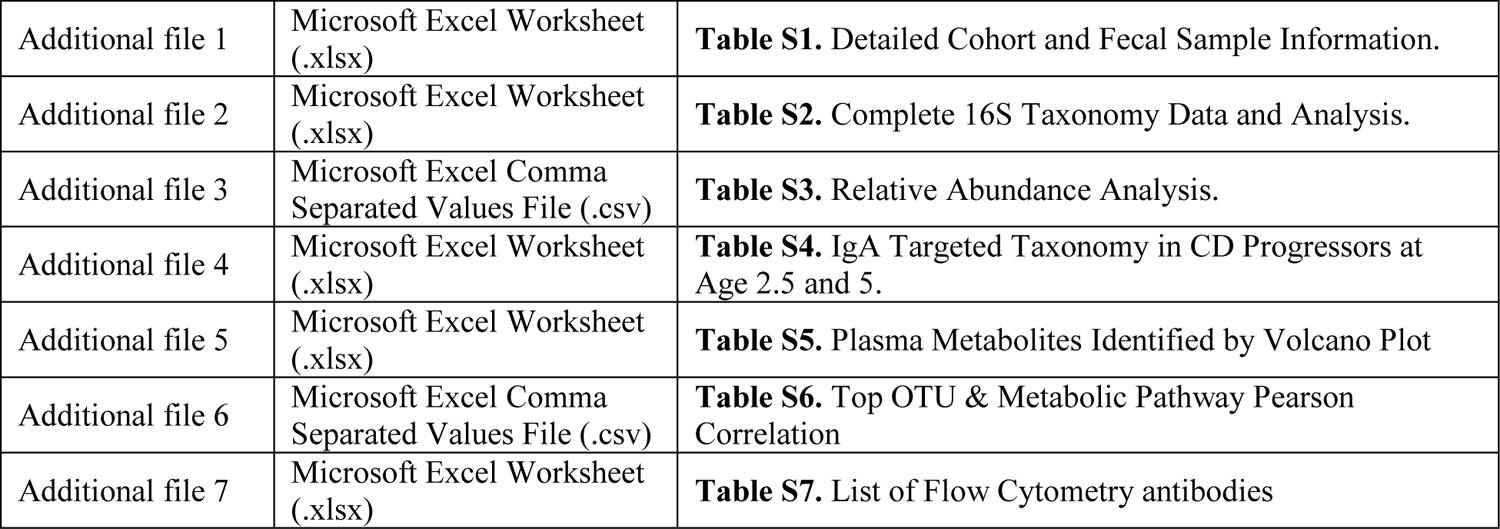

|  |  |  |
| --- | --- | --- |
| Additional file 1 | Microsoft Excel Worksheet (.xlsx) | **Table S1.** Detailed Cohort and Fecal Sample Information. |
| Additional file 2 | Microsoft Excel Worksheet (.xlsx) | **Table S2.** Complete 16S Taxonomy Data and Analysis. |
| Additional file 3 | Microsoft Excel Comma Separated Values File (.csv) | **Table S3.** Relative Abundance Analysis. |
| Additional file 4 | Microsoft Excel Worksheet (.xlsx) | **Table S4.** IgA Targeted Taxonomy in CD Progressors at Age 2.5 and 5. |
| Additional file 5 | Microsoft Excel Worksheet (.xlsx) | **Table S5.** Plasma Metabolites Identified by Volcano Plot |
| Additional file 6 | Microsoft Excel Comma Separated Values File (.csv) | **Table S6.** Top OTU & Metabolic Pathway Pearson Correlation |
| Additional file 7 | Microsoft Excel Worksheet (.xlsx) | **Table S7.** List of Flow Cytometry antibodies |

## Methods

### Human Fecal Samples

The fecal samples were obtained from subjects in the All Babies in Southeast Sweden (ABIS) cohort. The ABIS study was ethically approved by the Research Ethics Committees of the Faculty of Health Science at Linköping University, Sweden (Ref. 1997/96287 and 2003/03-092) and the Medical Faculty of Lund University, Sweden (Dnr 99227, Dnr 99321). All children born in southeast Sweden between 1^st^ October 1997 and 1^st^ October 1999 (n=21,700) were eligible to participate and offered to participate and 17,055 families participated. Among them, 181 children developed celiac disease. Informed consent from the parents was obtained after oral and written information and information via video film. Fresh fecal samples were collected either at home or at the clinic. The questionnaire was completed by the parents to collect participants’ health information including, but not limited to, breastfeeding duration, antibiotic use, gluten exposure time, and more. In total 68 fecal samples were collected for the analysis (32 at age 2.5 and 26 at age 5). Diagnosis of CD was confirmed at least twice according to the Swedish National Diagnosis Register (Ludvigsson et al., 2011). Until 2012, the CD was diagnosed by both serology and small intestinal biopsy. Both records were registered on the Swedish national inpatient register (Ludvigsson et al., 2011).

### IgA+ and IgA-Bacteria Separation

IgA+ and IgA-bacteria separation was performed as previously described (Palm et al., 2014). Briefly, frozen human fecal samples were placed in Fast Prep Lysing Matrix D with ceramic beads (MP Biomedicals) and incubated in 1ml Phosphate Buffered Saline (PBS) per 100mg samples on ice for 5 min for hydrating. This was followed by homogenization using bead beating for 7s (Minibeadbeater; Biospec) and then centrifuged 50xg for 10 min at 4°C to remove large plaques. Fecal bacteria in the supernatants were collected (200 μl/sample) and washed three times with 500 μl PBS containing 1% (w/v) Bovine Serum Albumin (BSA, American Bioanalytical) and centrifuged for 5 min (6,000 x rpm, 4°C). A sample of this washed bacterial suspension (50 μl) was collected as the pre-sorting sample for 16S sequencing analysis. After washing, bacterial pellets were re-suspended in 50 μl blocking buffer (PBS containing 1% (w/v) BSA and 20% Normal Mouse Serum (Jackson ImmunoResearch), incubated for 20 min on ice, and stained with 100 μl PE-conjugated mouse anti-human IgA (1:40; Miltenyi Biotec clone IS11-8E10) for 30 minutes on ice. Samples were subsequently washed three times with 500 μl BSA containing 1% (w/v) before flow cytometry analysis or cell separation. PE anti-human IgA-stained bacteria were incubated with Anti-PE Magnetic Activated Cell Sorting (MACS) beads (Miltenyi Biotec) (1:5) for 30 minutes on ice and then separated by a custom magnetic plate for 10 minutes on ice. Fecal bacteria bound to the magnetic plate were collected as IgA+ samples for 16S sequencing analysis. Stained and MACS bead-bound bacteria unbound to magnet plate were collected (20∼40 μl) and passed through MACS molecular columns (Miltenyi Biotec) (one sample/column) followed by flushing with 480 μl PBS containing 1% (w/v) BSA. The total pass-through (∼500 μl) was loaded onto columns one more time. The columns were flushed with 500 μl PBS containing 1% (w/v) BSA. The total column pass-through (∼1 ml) was saved as IgA-samples for 16S sequencing analysis.

### Fecal IgA Flow Cytometry Analysis

Bacterial cells were isolated from fecal samples as described in the IgA+ and IgA-bacteria separation method section of this manuscript. Bacteria were stained with PE Anti-human IgA antibodies (1:100; Miltenyi Biotec clone IS11-8E10) for 30 min on ice. After washing twice, bacteria were stained with TO-PRO®-3 (ThermoFisher Scientific) to identify bacteria from fecal debris or particles. Stained bacteria were analyzed by a BD FACSAria^TM^ IIIu cell sorter (Becton-Dickinson) as previously described (Wilmore et al., 2018) as TO-PRO®-3^+^IgA^+/-^ cells.

### 16S rRNA Gene Sequencing

16S rRNA sequencing of the V4 region sequencing for all bacterial samples were performed on the Miseq platform with barcoded primers. Briefly, all bacterial samples were suspended in 90 μl of MicroBead Lysis Solution with 10% RNAse-A, and sonicated in a water bath at 50°C for 5 minutes. Samples were transferred to a plate containing 50 μl of Lysing Matrix B (MP Biomedicals) and homogenized by bead-beating for 5 minutes. After centrifugation (4122 x *g*, 4°C) for 6 minutes, the supernatant was transferred to 2 ml deep-well plates (Axygen Scientific). Bacterial DNA from the samples was extracted and purified using MagAttract Microbial kit (QIAGEN) following the instruction provided by the manufacturer. PCR was performed to amplify the V4 region of 16S ribosomal RNA (33 cycles) in duplicate (3 μl purified DNA per reaction; Phusion DNA polymerase, New England Bioscience) (Palm et al., 2014). After amplification, PCR products were then normalized with SequalPrep^TM^ normalization plate kit (ThermoFisher Scientific) and pooled. The pooled library concentration was calculated by using the NGS Library Quantification Complete kit (Roche 07960204001) and then loaded on a Miseq sequencer. Illumina Miseq Reagent Kit V2 (500 cycles) was used to generate 2×250bp paired-end reads. The raw reads were demultiplexed in Qiime1 (version 1.9). The sequencing yielded a mean of 30,471 reads per sample.

### Bioinformatics Analysis and Statistics

Microbial diversity and statistical analyses were performed by filtering and trimming of the bacterial 16s rRNA amplicon sequencing reads, and sample inference that turns amplicon sequences into an Amplicon Sequence Variant (ASVs) table was performed by dada2 (Callahan et al., 2016) using the Ribosomal Database Project Training Set 16(Cole et al., 2014).

Exploratory and inferential analyses were performed in R (version 4.1.2) using phyloseq (McMurdie and Holmes, 2013) and vegan (*al*, 2019), which includes non-metric multidimensional scaling (NMDS) analysis using Bray–Curtis dissimilarity, Principle Components Analysis (PCA), alpha and beta diversity estimates, and taxa agglomeration. Statistical significance was assessed by ANOVA for alpha diversity and PERMANOVA for Bray-Curtis dissimilarity. Differential ASVs abundance was assessed per time point by edgeR (Robinson et al., 2010) with two-sided empirical Bayes quasi-likelihood F-tests. P-values were corrected by using the Benjamini-Hochberg false discovery rate (FDR), and FDR < 0.05 was considered a statistically significant (Benjamini, 1995).

### Plasma Metabolomics Analysis

Plasma samples for metabolomics analysis were prepared as previously described (Baskin et al., 2018; Drolet et al., 2017). Metabolite extraction from plasma was achieved using a mixture of isopropanol, acetonitrile, and water at a ratio of 3:3:2 v/v. Extracts were divided into three parts: 75 μl for gas chromatography combined with time-of-flight high-resolution mass spectrometry, 150 μl for reverse-phase liquid chromatography coupled with high-resolution mass spectrometry, and 150 μl for hydrophilic interaction chromatography with liquid chromatography and tandem mass-spectrometry, and analyzed as previously described (Baskin et al., 2018; Drolet et al., 2017). We used the NEXERA XR UPLC system (Shimadzu, Columbia, MD, USA) coupled with the Triple Quad 5500 System (AB Sciex, Framingham, MA, USA) to perform hydrophilic interaction liquid chromatography analysis, the NEXERA XR UPLC system (Shimadzu, Columbia, MD, USA) coupled with the Triple TOF 6500 System (AB Sciex, Framingham, MA, USA) to perform reverse-phase liquid chromatography analysis, and an Agilent 7890B gas chromatograph (Agilent, Palo Alto, CA, USA) interfaced to a Time-of-Flight Pegasus HT Mass Spectrometer (Leco, St. Joseph, MI, USA). The GC system was fitted with a Gerstel temperature-programmed injector cooled injection system (model CIS 4). An automated liner exchange (ALEX) (Gerstel, Muhlheim an der Ruhr, Germany) was used to eliminate cross-contamination from the sample matrix that was occurring between sample runs. Quality control was performed using metabolite standards, mixture, and pooled samples. A standard quality control sample containing a mixture of amino and organic acids was injected daily to monitor mass spectrometer response. A pooled quality control sample was obtained by taking an aliquot of the same volume from all samples of the study and injecting daily with a batch of analyzed samples to determine the optimal dilution of the batch samples and validate metabolite identification and peak integration. Collected raw data were manually inspected, merged, inputted, and normalized by the sample median. Metabolite identification was performed using in house authentic standards analysis. Metabolite annotation was used utilizing recorded retention time and retention indexes, recorded MS^n^ and HRAMS^n^ data matching with METLIN, NIST MS, Wiley Registry of Mass Spectral Data, HMDB, MassBank of North America, MassBank Europe, Golm Metabolome Database, SCIEX Accurate Mass Metabolite Spectral Library, MzCloud, and IDEOM databases.

### Metabolite pathway analysis

Metabolomic data were analyzed as previously described by Tolstikov et al. (Tolstikov et al., 2014). Identified metabolites were subjected to pathway analysis with MetaboAnalyst 4.0, using a Metabolite Set Enrichment Analysis (MSEA) module which consists of an enrichment analysis relying on measured levels of metabolites and pathway topology and provides visualization of the identified metabolic pathways. Accession numbers of detected metabolites (HMDB, PubChem, and KEGG Identifiers) were generated, manually inspected, and utilized to map the canonical pathways. MSEA was used to interrogate functional relation, which describes the correlation between compound concentration profiles and clinical outcomes.

### Animals

C57BL/6J mice were maintained and bred in the Boston College Animal Care Facility. All the animal experiments were conducted following the regulations and ethics guidelines of the National Institute of Health and were approved by the IACUC of Boston College (Protocol No.#B2019-003 and 2019-004). The mice were maintained under specific pathogen-free conditions in a 12-hrs dark/light cycle with free access to autoclaved water, and bedding. After weaning, 3-week-old mice were divided into two groups: (1) Control group, were provided with normal diet and, (2) TDCA group, were provided with normal diet containing 0.4% TDCA. Mice were maintained on their diets for up to 10 weeks. After 10 subsequent weeks of diet, mice were sacrificed and the primary cell suspension was isolated from mesenteric lymph nodes (MLNs), intraepithelial cells (IELs), Peyer’s patches (PPs), and Lamina propria (LPs) and spleen were isolated

### Primary cell isolation and Flow cytometry

Cells were isolated from the spleen and MLN via mechanical disruption, and LPs and PPs were isolated as described previously. Briefly, Intestines were washed with PBS and different parts were cut transversally into small 1cm pieces and fixed in formalin. The intestines were then cut longitudinally to expose the inner epithelium layer and were washed with PBS 2-3 times to get rid of feces containing fetal bovine serum (FBS). The cut pieces of intestines were stirred in freshly prepared dithiothreitol (DTE) solution (HBSS (1x), HEPES-bicarbonate solution (1x), containing 1 mmol/l DTE supplemented with 10% FBS (Gibco) for 20 mins two times. After 20 mins, IELs were collected from the supernatant and filter using a 70μm filter and then passed through a 44/67 percoll gradient. The intestine pieces were stirred in EDTA solution (i.e HBSS, HEPES-bicarbonate solution containing 1.3 mmol/l EDTA, 2mM glutamine and antibiotics (Gibco)) for 30 mins a total of two times and then washed with complete media. To isolate LPs and Peyer’s patches (PPs), the washed intestines and PPs were resuspended in collagenase solution (RPMI-1640 medium (Sigma-Aldrich) containing HEPES solution, (1x) 1mM of MgCl_2_ and 1mM of CaCl_2_ supplemented with 10% FBS, 2 mmol/l L-glutamine, antibiotics and collagenase (100 U/mL) were collected from the supernatant by passing through 70 μm filter and then passed through a 44/67 percoll gradient. Cells were isolated from the gradient, centrifuge, washed, and filtered to get a single-cell suspension. The cells were stained with trypan blue and live cells were counted manually using a hemocytometer. The cell suspension was then washed surface labeled with the appropriate fluorochrome-conjugated monoclonal antibodies as mentioned in **Table S6**. The results were assessed using flowjo10 software.

### Histopathological sectioning and staining

Structural and cellular alteration induced by TDCA in the intestines of the mice was determined by histology. After ten weeks of TDCA treatment, the different parts of the small intestine (duodenum, jejunum, ileum) and colon were washed with PBS and fixed in 10% formalin (v/v) at 4°C overnight and washed and stored in 70% ethanol. The parafilm blocks of ileum were prepared, and 5 μm thick sections were cut using a microtome (Leica). The ileum sections were dried, stained with hematoxylin and eosin (H&E) staining kit (Vector Laboratory), and mounted using permount (Fisher scientific). The sections were analyzed using the upright microscope (Zeiss AxioImager Z2). The quantification of the length of villi, crypt, and the number of cells was performed using Fiji/ ImageJ software.

---

**Figure S1.**
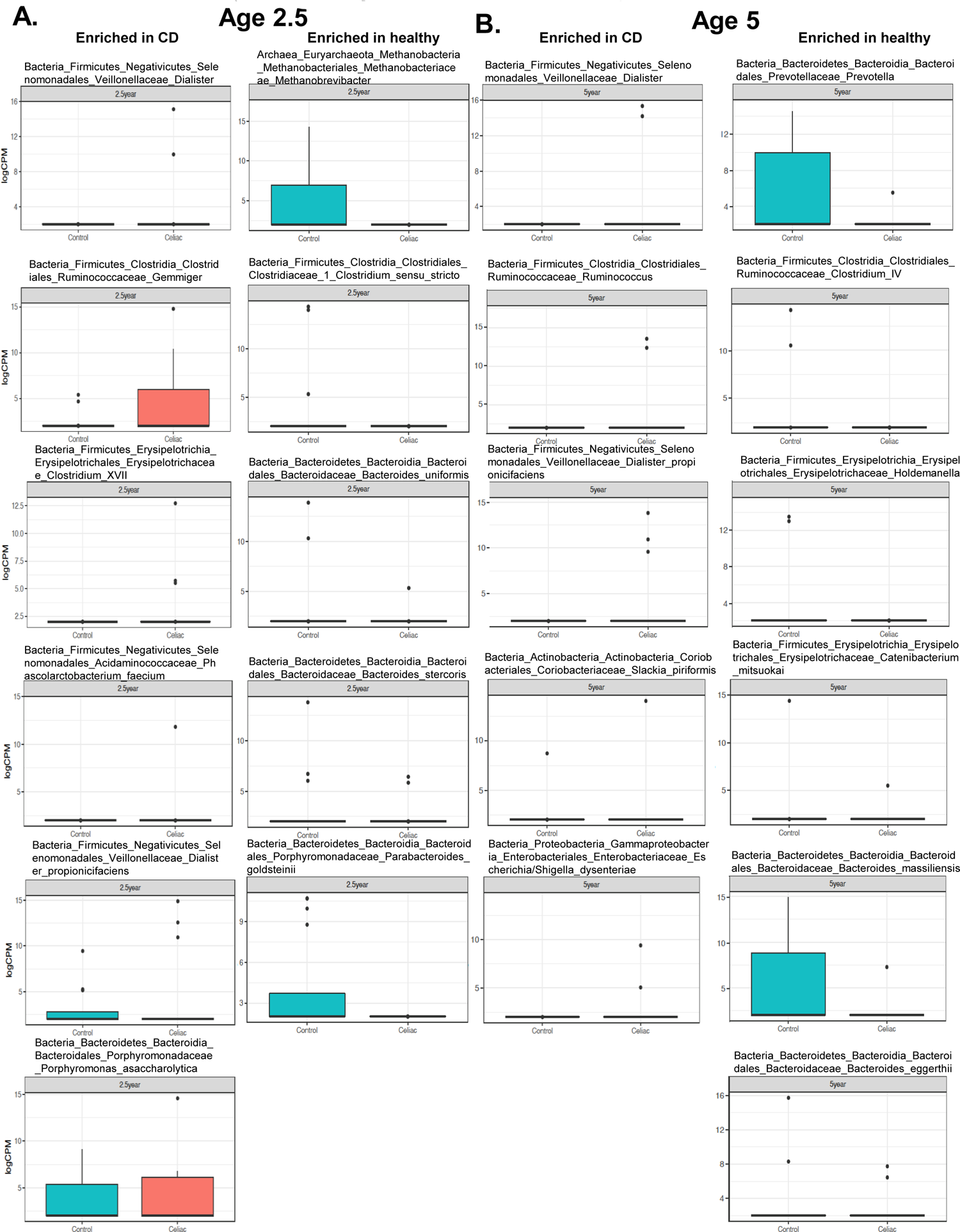
Box plot representation of most enriched genus/species in CD progressors compared to healthy controls at ages 2.5 and 5. A. Fold change in ASVs at age 2.5 in CD progressors (left panel) and healthy controls (right panel) B. Fold change in ASVs at age 5 in CD progressors (left panel) and healthy controls (right panel).

**Figure S2.**
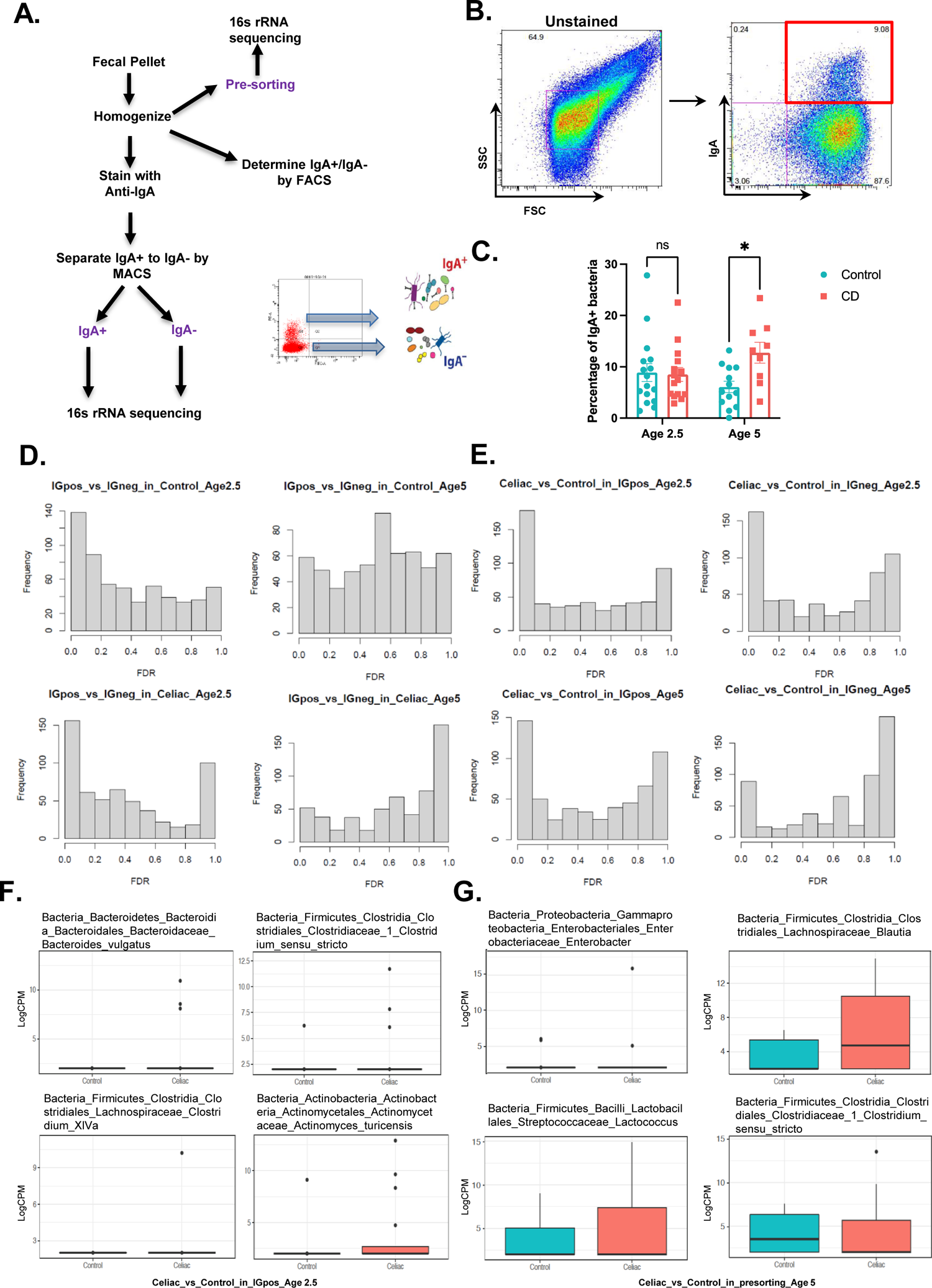
Gating strategy for IgA sequencing and analysis. **A.** Schematic overview of IgA-based fecal bacteria separation combined with 16S rRNA gene sequencing (IgA-seq) for stool samples from CD progressors and healthy controls. MACS: Magnetic-activated cell separation. **B.** Gating strategy for the isolation of IgA-/+ bacteria from the CD progressors and healthy controls’ fecal samples. **C**. Flow cytometry results for IgA coated (IgA+) bacteria from CD progressors and healthy controls at age 2.5 and 5. Three CD progressors who were diagnosed with CD diagnosis before 5 were excluded in this analysis. (age 2.5, n=15-16; and age 5: n=9-13). Data were expressed as mean ± SEM. *P<0.05, **P<0.01, ***P<0.01. Statistical analysis was performed by two-way ANOVA. **D.** Empirical Bayes quasi-likelihood F-tests analysis for the comparisons of IgA coated and non-coated gut microbiota ASVs in healthy controls (upper row) and CD progressors (lower row) at ages 2.5, and 5. Frequency: number of ASVs. FDR: False Discovery Rate. **E.** Empirical Bayes quasi-likelihood F-tests analysis for the comparisons of IgA coated or non-coated gut microbiota ASVs between CD progressors and healthy controls (upper row: age 2.5 years old; lower row: age 5 years old). **F.** Box plots showing representative ASVs in which abundances were similar in the gut microbiota (presorting samples) but differently targeted by IgA at age 2.5. **G.** Box plots showing representative ASVs in which abundances were similar in the gut microbiota (presorting samples) but differently targeted by IgA at age 5.

**Figure S3.**
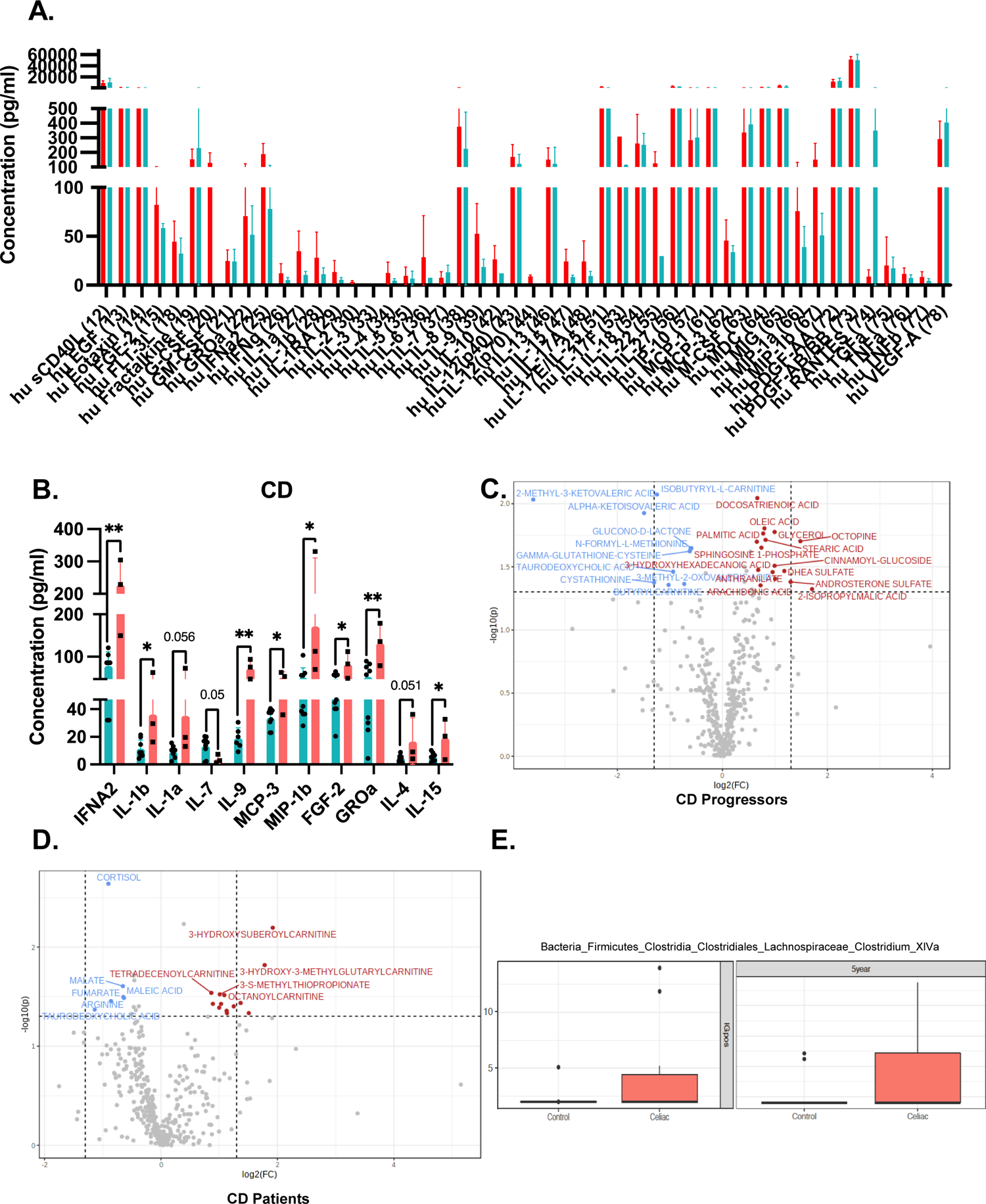
The volcano plot of plasma metabolome of CD progressors and CD patients. **A.** Comparison of all 48 cytokines analyzed in plasma samples obtained from CD progressors (n=10) and healthy controls (n=10) at age 5. Data were expressed as means ±SEM. *, P<0.05, **, P<0.01, ***, **B.** Significantly altered cytokines in only CD patients (n=3) in comparison to healthy controls (n=10). Data were expressed as means ±SEM. *, P<0.05, **, P<0.01, ***, P<0.001. Statistical analysis was performed by a two-tailed, unpaired Student’s t-test. **C.** Volcano plot of plasma metabolites of CD progressors (n=7) -excluding patients who develop CD within 5yrs (n=3) vs. healthy controls (n=9) with fold change threshold (|log_2_ (FC)|>1.2) and t-tests threshold (-log_10_(p)>0.1). Colored dots represent metabolites above the threshold. Fold changes are log_2_ transformed, and p values are log_10_ transformed. **D.** Volcano plot of plasma metabolites of CD patients (n=3) who developed CD before age 5 vs. healthy controls (n=9) with fold change threshold (|log_2_ (FC)|>1.2) and t-tests threshold (-log_10_(p)>0.1). The colored dots represent metabolites above the threshold. Fold changes are log_2_ transformed, and p values are log_10_ transformed. **E.** Box plots showing the representative *Clostridium XIVa* bacteria abundance between CD progressors and healthy controls (Left: before separation by IgA coating, Right: in IgA+ bacteria).

**Figure S4.**
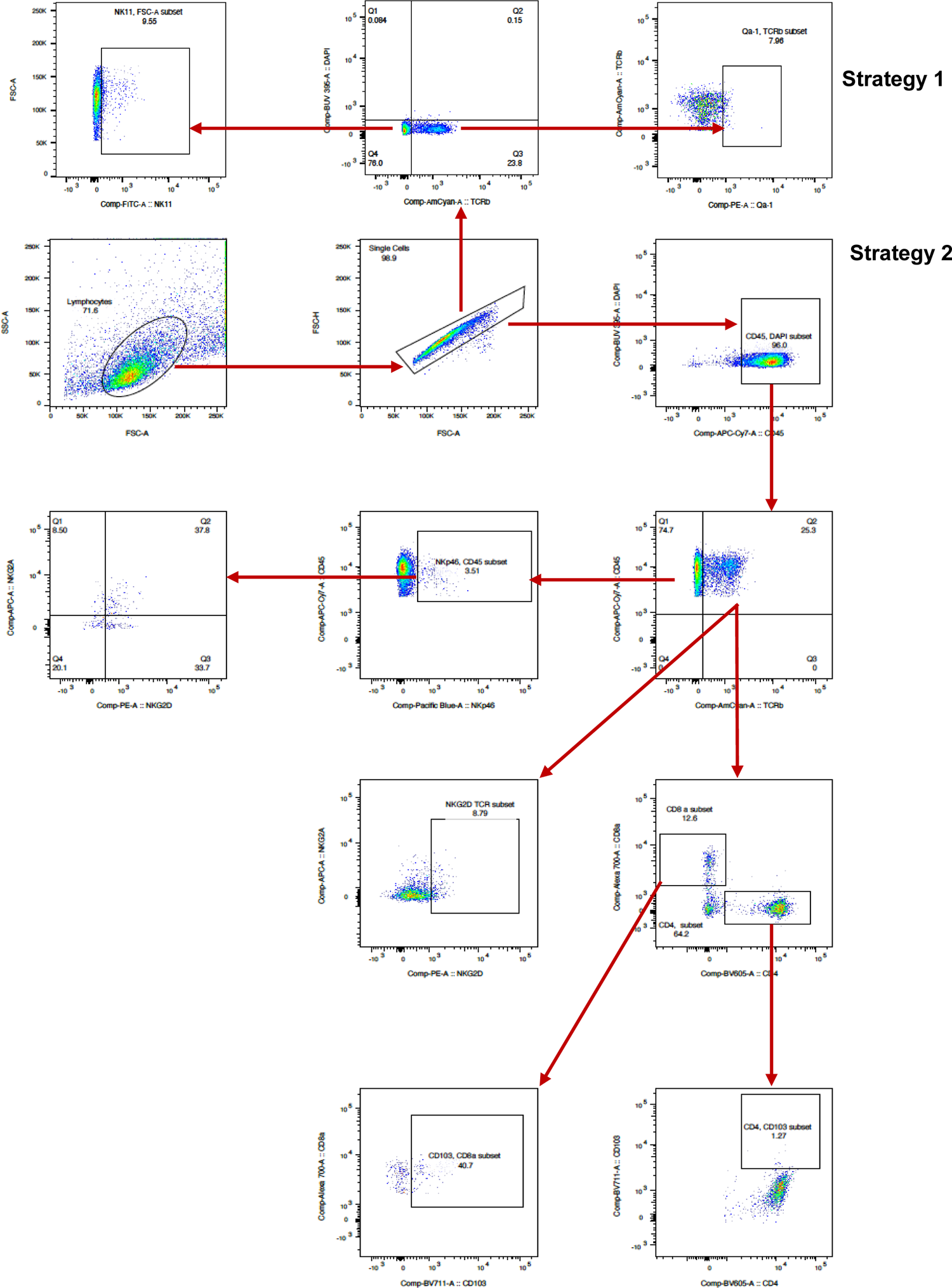
Gating Strategy for the Flow cytometry analysis. Strategy 1-Describing the gating strategy for NK1.1 and Qa-1 expression in TCRβ+ cells. Strategy 2-Describing the gating strategy for CD8, CD4, NKG2D, CD103, and NKp46.

**Figure S5.**
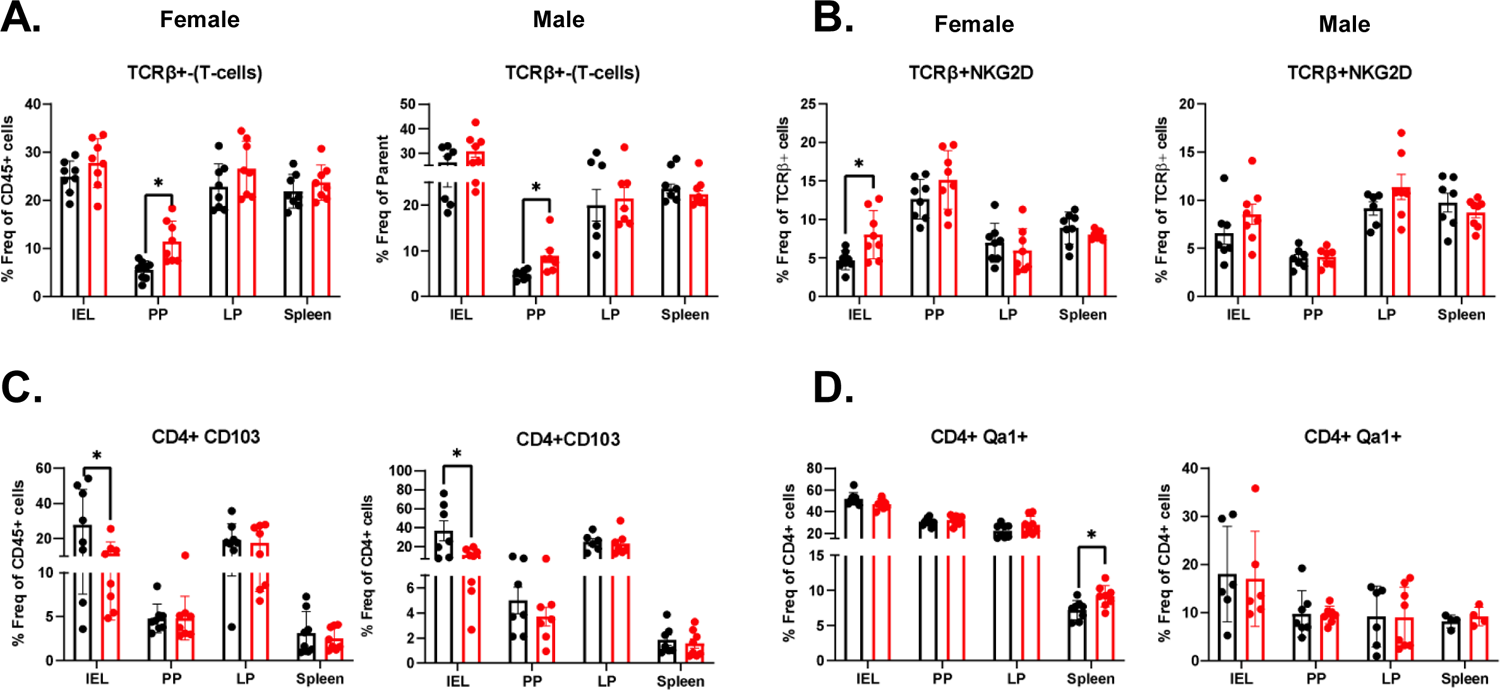
TDCA diet induces changes in T-cell composition in different cell subsets. **A.** TCRβ+ cells as % of total CD45+ cells in the IELs, PP, LP and spleen of female (left panel) and male (right panel) mice. **B.** NKG2D+ cells as % of total TCRβ+ CD45+ cells in the IELs, PP, LP and spleen of female (left panel) and male (right panel) mice. **C.** CD103+ cells as % of total CD4+ cells in the IELs, PP, LP and spleen of female (left panel) and male (right panel) mice. **D.** Qa-1+ cells as % of total CD4+ cells in the IELs, PP, LP and spleen of female (left panel) and male (right panel) mice. Data were expressed as mean ± SEM. *P<0.05, **P<0.01, ***P<0.01. Statistical analysis was performed by a two-tailed unpaired Student’s t-test.

